# Using resurvey data to predict changes in ecosystem functioning across protected and unprotected coastal dunes

**DOI:** 10.1101/2025.03.31.646285

**Authors:** Greta La Bella, Alicia T.R. Acosta, Tommaso Jucker, Manuele Bazzichetto, Marco Andrello, Marta Gaia Sperandii, Marta Carboni

## Abstract

Protected areas are generally designed to conserve biodiversity. However, how well they also contribute to maintaining ecosystem functions that plant diversity supports has rarely been explicitly tested, often due to the lack of historical ecosystem function data. Here, we used a trait-based approach to reconstruct past ecosystem functioning and examine its change over the last 15 years in protected and unprotected coastal dune ecosystems, checking where functions remain stable over time. First, we resurveyed vegetation in quasi-permanent plots and measured in the present several ecosystem functions related to biomass production, carbon, water, nutrient cycling, erosion control, and invasion resistance across six coastal dune sites in Central Italy. Second, using these data, we quantified Biodiversity-Ecosystem Function (BEF) relationships and employed them to hindcast past ecosystem functions based on historical vegetation surveys. Finally, as a case study, we applied this method to assess temporal changes in ecosystem functioning under three protection regimes: national protected areas (i.e. strict protection), Natura 2000 sites (loose protection), and non-protected areas. Biomass production, carbon, and water regulation increased over time in non-protected areas, likely due to an expansion of ruderal and non-native species, that are usually more productive. Within Natura 2000 sites, communities showed a decrease in erosion control and invasion resistance, due to the loss of important dune-building species and the spread of non-natives. Only within national protected areas, ecosystem functions did not undergo significant temporal changes, and invasion resistance even increased. Our results suggest that ecosystem functioning remained stable over time only in areas under strict protection. More broadly, our study demonstrates the potential for using revisitation data in combination with locally estimated BEF relationships to hindcast past ecosystem functioning, providing a valuable tool for monitoring long-term functional changes in response to conservation measures.

## INTRODUCTION

Monitoring ecosystem functions is critical for assessing the health and resilience of ecosystems and ensuring their ability to provide essential services to human societies. The post-2020 Global Biodiversity Framework (GBF) of the Convention on Biological Diversity (CBD) emphasizes not only the need to halt biodiversity loss but also to maintain and enhance ecosystem functions and services, as reflected in its goals and targets (CBD, 2022). Similarly, the United Nations Sustainable Development Goals (SDGs) highlight the importance of sustaining ecosystem integrity to support human well-being (UN, 2015). Despite these global commitments, measuring ecosystem functions presents significant challenges due to their complex, multi-dimensional nature and the difficulty of obtaining long-term data. Many ecosystem processes operate at large spatial and temporal scales, requiring indirect methods or models to infer past dynamics. This limitation complicates the evaluation of conservation strategies, such as protected areas and restoration efforts, which should aim to preserve not only species diversity but also the ecological processes that sustain ecosystems. Without robust estimates of past ecosystem functions, it remains difficult to determine whether conservation actions have successfully maintained or restored functional integrity.

The relationship between Biodiversity and Ecosystem Functioning (BEF) has been widely documented across natural ecosystems, although its strength and direction can vary depending on the local environmental conditions and community composition (Hagan et al., 2021; Hanisch et al., 2020; La Bella et al., 2024; Ratcliffe et al., 2017; Valencia et al., 2015; van der Plas, 2019). Given this relationship, shifts in biodiversity can significantly alter ecosystem processes and functions. Nevertheless, while biodiversity changes are increasingly being reported, only few studies have quantified the extent and direction in which ecosystem functions are changing (Brauman et al., 2020). As ecosystem functions largely underpin vital services provided to people (Garland et al., 2021; MEA, 2005), monitoring changes in ecosystem functioning is of crucial importance for human well-being.

To prevent further biodiversity loss and safeguard the last remaining intact natural areas, European countries have implemented national or international conservation measures (Heslenfeld et al., 2008), including the establishment of protected areas (CBD, 2022; UN, 2015). By conserving biodiversity protected areas also offer the opportunity to preserve the variety of ecosystem functions and processes that biodiversity supports. However, since most protected areas are not explicitly managed to maintain ecosystem functionality, their ability to counteract changes in ecosystem functions remains uncertain.

Overall, studies assessing temporal changes in ecosystem functioning in real-world ecosystems are scarce for several reasons (DeCock et al., 2023; Waldén et al., 2023; Williams et al., 2005). First, the interest in quantifying ecosystem functions in natural systems has only surged in recent years (Garland et al., 2021; van der Plas, 2019). Although systematic data collection has been established in some long-term ecological monitoring networks (e.g. the Biodiversity Exploratories; Fischer et al., 2010), these remain relatively localised. As a result, comparison with past conditions are still limited (Kremen & Ostfeld, 2005; Meyer et al., 2015). Second, unlike biodiversity metrics, measuring ecosystem functions is challenging and methods are still highly debated (Byrnes et al., 2014; Garland et al., 2021; Manning et al., 2018; Meyer et al., 2015). In fact, while some ecosystem functions can be quantified through direct, well-accepted indicators (e.g. biomass production), for other functions choosing the most appropriate indicator is less clear (e.g. water regulation) (Farnsworth et al., 2017; Garland et al., 2021; Jax, 2005). Moreover, measuring ecosystem functions is usually resource-intensive and time-consuming, requiring technical expertise and equipment which were until recently not widely accessible (Meyer et al., 2015). Consequently, reconstructing past data in order to track temporal changes in ecosystem functioning requires innovative solutions.

Plant functional traits are increasingly recognised as reliable predictors of several ecosystem functions and processes, such as biomass production, carbon sequestration, and nutrient cycling (de Bello et al., 2010; Díaz et al., 2007; Funk et al., 2017; Lavorel & Garnier, 2002). As a result, plant traits are now routinely used in BEF research together with taxonomic diversity for quantifying the effect of species and communities on ecosystems. For example, species-rich communities dominated by fast-growing acquisitive plant species have been frequently associated to fast biogeochemical cycles (Cornwell et al., 2008; Funk et al., 2017; Garnier et al., 2015; Lavorel & Garnier, 2002). Given this relationship, several trait-based approaches have been proposed to predict ecosystem functioning beyond the observed data, e.g. to larger datasets (Bodenhausen et al., 2023; Chen et al., 2023), or for predicting ecosystem functions under future scenarios of environmental change (Klumpp & Soussana, 2009; Madani et al., 2018; Quétier et al., 2007). BEF relationships can thus potentially also be used to hindcast past functioning levels using historical data, with even less uncertainty compared to e.g. forecasts based on many possible future scenarios. This potential application provides a chance to compare past and present ecosystem functions and, thus, to assess how ecosystems have responded to the last decades of global changes. As such, here we propose predictive modelling to reconstruct past ecosystem functions based on historical vegetation composition in order to track temporal changes in functioning, particularly in highly endangered ecosystems.

Coastal dunes are very peculiar ecosystems, rich in highly specialized flora with an outstanding conservation value (Acosta et al., 2009; Van der Maarel, 2003). They also provide multiple essential services to society, e.g. coastal protection against the sea (Liquete et al., 2013) and recreational activities (Drius et al., 2019a). Despite their valuable features, coastal dunes are among the most threatened ecosystems in Europe, especially in the Mediterranean basin (Janssen et al., 2016; Prisco et al., 2020). High anthropogenic impact related to urban expansion and tourism as well as the spread of invasive alien species, have led to extensive degradation of coastal habitats and to biodiversity declines (Carboni et al., 2010; Defeo et al., 2009; Drius, Bongiorni, et al., 2019; Malavasi et al., 2016; Schlacher et al., 2007). Still, the consequences of habitat degradation for the functioning of coastal dune ecosystems have been rarely quantified.

In this study, we tackled two key objectives: (1) testing the predictive ability of vegetation resurveys for hindcasting ecosystem functioning and assessing possible gains and losses in functions; (2) exploring the performance of different types of protected areas in maintaining the level of ecosystem functioning stable over time. Specifically, we developed an approach that integrates vegetation resurvey data with biodiversity–ecosystem functioning (BEF) models to explore temporal trends in key ecosystem functions. We then applied this framework (Fig. 1) to coastal dune ecosystems, comparing temporal changes in ecosystem functioning across three protection regimes: nationally designated strictly protected areas, sites belonging to the European Natura 2000 network (protected areas that allow a sustainable use of natural resources), and non-protected areas.

**Figure 1.**
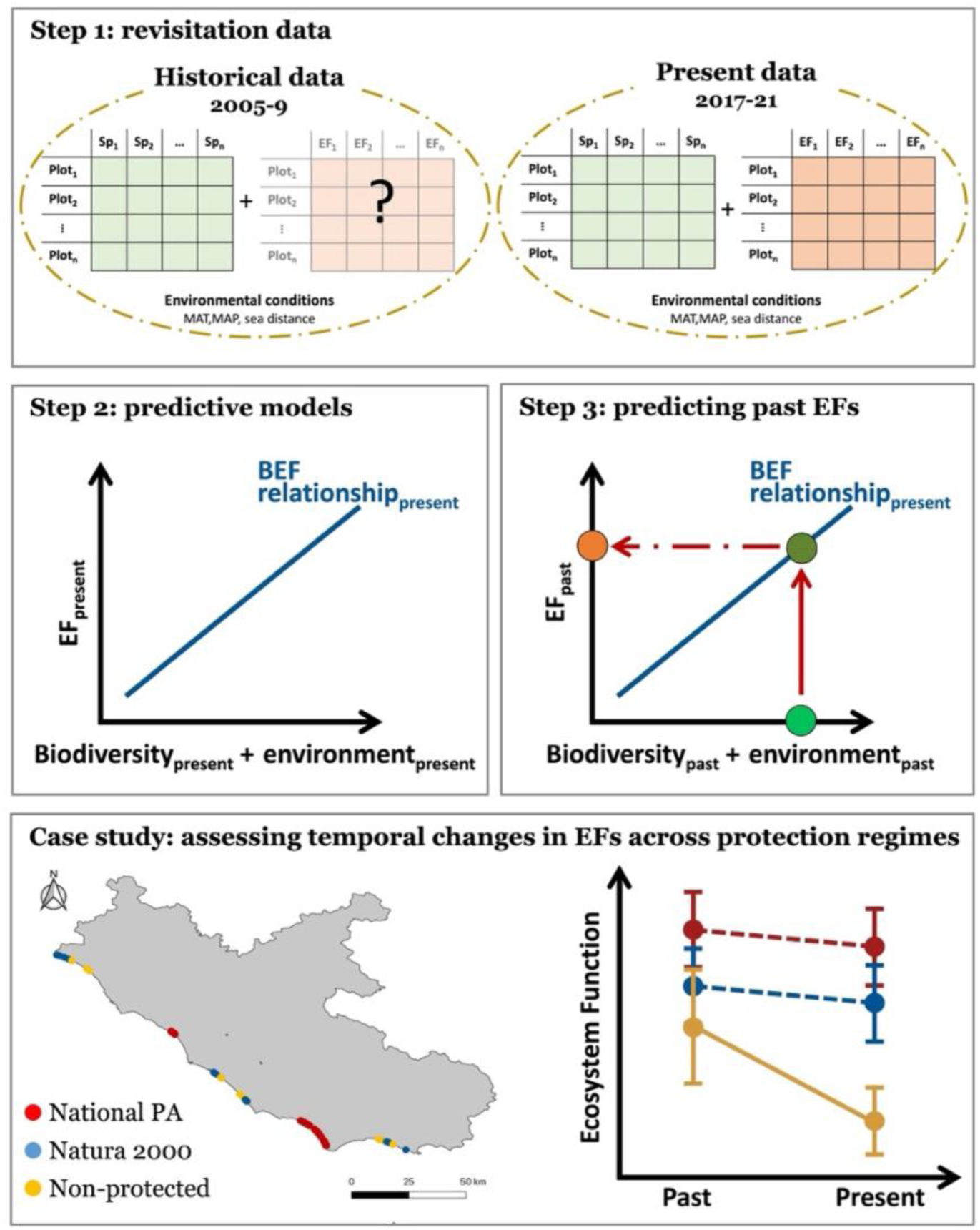
Workflow used to assess temporal changes in ecosystem functioning. **Step 1**: in 2021, we resurveyed vegetation in quasi-permanent plots and collected several ecosystem function variables. **Step 2**: using these data, we assessed BEF relationships. **Step 3**: we employed BEF relationships to hindcast past ecosystem functioning based on historical vegetation surveys. **Case study:** we used this approach to assess temporal changes in ecosystem functioning across different protection regimes, i.e. national protected areas (PA), Natura 2000 sites, and non-protected areas, along the coastal dunes of Lazio region (Italy).

## METHODS

### Study area and protection status

The research was conducted on coastal dune ecosystems situated along the Tyrrhenian coast of Central Italy, specifically in the region Lazio, consisting of approximately 250 km of coast. This geographical area falls within the Mediterranean climatic region and consists of Holocene sandy dunes forming a narrow stretch along the shoreline (Acosta et al., 2009).

For determining the protection status, we acquired spatial data related to the World Database on Protected Areas (WDPA) and World Database on Other Effective Area-based Conservation Measures (WD-OECM), from the “Protected Planet” data portal (UNEP-WCMC and IUCN, 2025). We considered three categories of protection: nationally designated protected areas, including both national parks and natural reserves (hereafter national protected areas); sites belonging to the Natura 2000 network, under Habitat Directive 92/43/EEC and Bird Directive 79/409/EC, which, compared to national protected areas allow a sustainable, albeit limited, use of nature (e.g., controlled dune stabilization or regulated tourism; Dudley, 2008; EC, 2025; Tsiafouli et al., 2013); and areas without any legal protection status. In cases of overlap between protection categories, such as Natura 2000 sites located within national protected areas, we assigned the highest protection level.

### Historical and revisitation data

We used an Italian database of coastal dune vegetation resurveying (“RanVegDunes” dataset; Sperandii et al., 2017) consisting for the study area in 286 quasi-permanent random plots (4 m^2^), i.e. georeferenced plots that can be reliably relocated over time, despite not being physically marked in the field (Kapfer et al., 2017).These plots were uniformly distributed across six study sites (Appendix S1) covering all herbaceous habitats of the coastal vegetation zonation along the sea-inland gradient in central Italy, i.e. the upper beach, embryo dunes, shifting dunes, and dune grasslands (see Acosta et al., 2007 for a description of the habitats). Overall, plot fall within two national protected areas, five Natura 2000 sites, and six non protected areas (Appendix S1 and S2). Plots were historically surveyed between 2005-2009 (historical data) and then revisited between 2017-2021 (revisitation data), thus after a period of 10-15 years. In each plot, sampling consisted in recording the number and percentage cover of all vascular plant species during the peak of the growing season (between April and May). Sampling method was consistent between historical survey and revisitation (see Sperandii et al., 2019 for details about the resurveying protocol). We classified species into natives and non-natives following (Galasso et al., 2024).

### Plant traits

We compiled functional trait data from local databases incorporated into TRY (Kattge et al., 2020), aggregating plant traits measured within the study area of Central Italy. Specifically, we chose four aboveground and four belowground traits: plant height (H; measured in m), leaf area (LA; measured in cm2), specific leaf area (SLA; measured in mm2/mg), leaf dry matter content (LDMC, measured in mg/g), root diameter (RD; measured in mm), specific root length (SRL; measured in m/g), root tissue density (RTD; measured in g/cm3), and root dry matter content (RDMC; measured in g/g). These traits are associated with the leaf and roots economic spectra (Carmona et al., 2021; Díaz et al., 2016), and are known to be related to several ecosystem processes and functions (Bardgett et al., 2014; de Bello et al., 2010; Freschet et al., 2021; Lavorel & Garnier, 2002; Roumet et al., 2016), including on coastal dunes (La Bella et al., 2023, 2024). Trait values were averaged across individuals by species, then log-transformed, centred, and scaled (Májeková et al., 2016).

### Environmental conditions

To account for climatic variability potentially affecting changes in vegetation between the two sampling periods, we obtained mean annual temperature (MAT), mean temperature in the coldest quarter (Bio11), mean annual precipitation (MAP), and precipitation in the warmest quarter (Bio18) of each plot from CHELSA, a high-resolution (30 arcs, ∼1 km) global database of climate data (Karger et al., 2021). However, due to high collinearity among these variables (see Appendix S3), we retained MAT as the sole climatic predictor in the models to avoid multicollinearity issues and improve model interpretability. In coastal dunes, the distance from the sea is commonly associated with a spatial gradient of environmental disturbance and, to a lesser extent, stress (Carboni et al., 2011). This means that communities located closer to the shoreline are typically exposed to higher levels of natural disturbances, such as tidal storms, salt spray, wind, and soil burial. Stress factors, such as water and nutrient limitation due to the high permeability of sandy soils, may also covary with sea distance, although not always linearly (McLachlan & Defeo, 2017). Despite such heterogeneity, sea distance remains a widely used proxy for capturing the composite sea-inland environmental gradient in coastal dune systems. Therefore, we included the shortest distance between each plot and the shoreline (hereafter sea distance) as a predictor in the models (Bazzichetto et al., 2016; La Bella et al., 2023). The shorelines of 2006 and 2017 were obtained through photointerpretation from satellite images in a QGIS environment (Version 3.28). We used the shoreline of 2006 for historical plots and the shoreline of 2017 for revisitation plots to quantify sea distance as the shortest Euclidean distance of each plot from the shoreline. As tidal variations along Mediterranean Sea are relatively small, ranging between 0.2-1 m (Fenu et al., 2013), the effect of tides on sea proximity measurement is negligible.

### BEF subset

Given the logistical constraints of measuring ecosystem functions across the entire dataset, we extracted from the revisitation dataset a subset of 110 plots to collect data related to ecosystem functioning in addition to plant biodiversity (hereafter, BEF subset). We used a stratified random approach based on biodiversity data and environmental conditions in order to represent the full range of variation observed across both historical and revisitation datasets. For this subset, in spring-summer season of 2021, we repeated the floristic survey (as explained above) and concurrently collected ecosystem function variables related to supporting and regulating services (Garland et al., 2021; MEA, 2005). Supporting services were biomass production, soil water regulation, soil carbon stock, and soil nutrients cycling, while as regulating services we assessed erosion control and invasion resistance (Table 1). Aboveground living plant biomass was used as an indicator of the total biomass produced by each plant community. While it does not represent a direct measure of net primary productivity, it reflects the production potential across plots (Allan et al., 2015; Cardinale et al., 2013; Garland et al., 2021; Ratcliffe et al., 2017; Sala & Austin, 2000). Water regulation, carbon stock, and nutrient cycling were quantified from soil samples collected in each plot. A full description of the sampling and analysis of these ecosystem functions can be found in Supporting Information (Appendix S4).

**Table 1.**
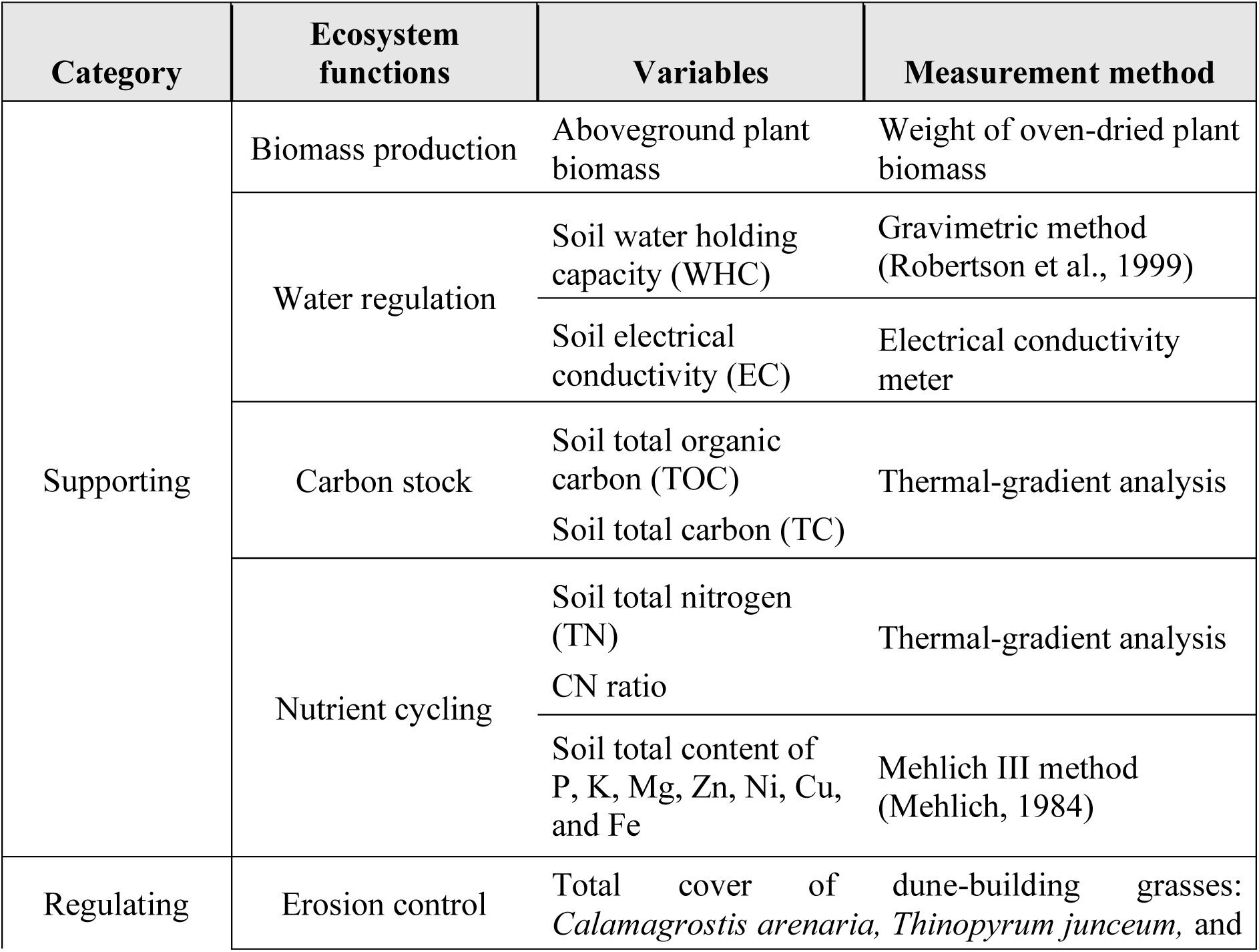
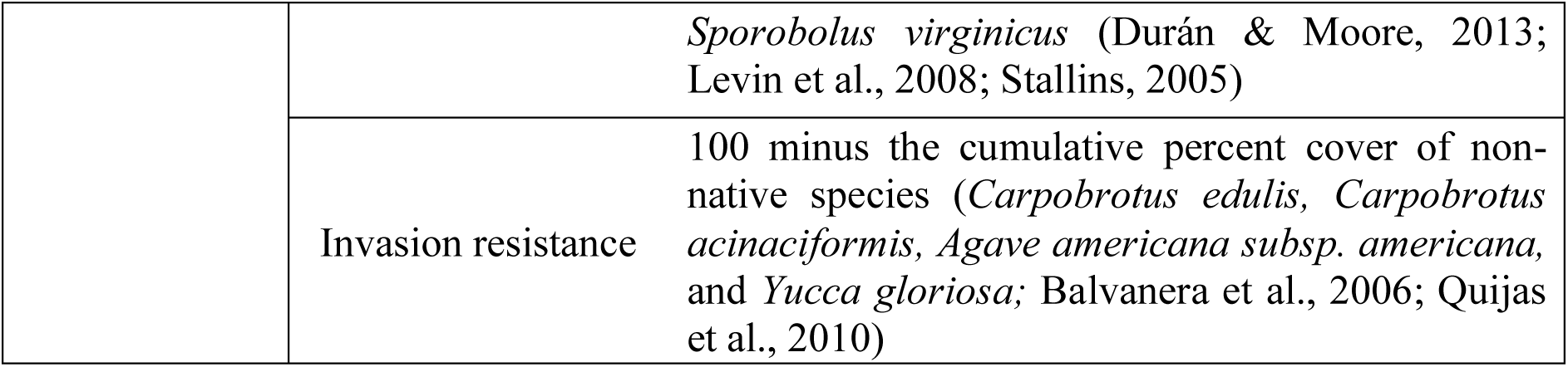
Variables and methods used to measure ecosystem functions.

In coastal dunes, erosion control is mainly operated by rhizomatous geophyte grasses that, through a dense root network, entrap sandy sediments and stabilize the dune. For this function, rhizomatous geophyte grasses are commonly referred to as dune-building or engineering species (Durán & Moore, 2013; Levin et al., 2008; Stallins, 2005). Here we used the total cumulative cover of the dune-building grasses *Calamagrostis arenaria, Thinopyrum junceum,* and *Sporobolus virginicus*, as a measure of the erosion control function. Invasion resistance was assessed as 100% minus the cumulative percent cover of the most abundant non-native species in the study area (Cascone et al., 2021), i.e. *Carpobrotus edulis*, *Carpobrotus acinaciformis*, *Agave americana subsp. americana*, and *Yucca gloriosa* (Balvanera et al., 2006; Quijas et al., 2010). Thus, higher values indicate greater resistance to invasion due to the lower cover of non-natives, while lower values indicate lower resistance to invasion.

### Biodiversity and ecosystem functioning metrics calculation

Our approach assumes that past trait-environment relationships have remained stable over time, but this may not always hold, particularly in ecosystems that have undergone significant anthropogenic or climatic changes. Consequently, although the original resurveying database included 286 plots (each visited twice), we only hindcasted ecosystem functions in plots where environmental conditions were expected to have remained relatively stable over time, minimizing the risk of violating this assumption. In particular, we excluded plots: (1) in which vegetation disappeared at the time of the resurvey due to anthropogenization or coastal erosion (n = 50); (2) belonging to the drift lines’ annual vegetation (n = 21), due to their ephemeral nature (Maun, 2009); (3) where species with trait information did not account for at least 80 % of the total vegetation (n = 8) (Pakeman & Quested, 2007); (4) where environmental conditions and biodiversity measures were out of the prediction range to avoid extrapolation (n = 25). Although plots where vegetation disappeared may provide interesting insights over the consequences of habitat loss, we chose not to consider them in the analysis due to the unfeasibility of assigning reliable values of ecosystem functions. The final dataset consisted of 203 plots with two repeated measures each, located either inside national protected areas (n = 69) and Natura 2000 sites (n = 89), or in non-protected areas (n = 45).

#### Biodiversity metrics

In each plot, we calculated several diversity metrics for historical and revisitation data. In particular, we assessed both taxonomic and functional aspects of the community, including species richness, total vegetation cover, above- and below-ground functional diversity (FD_above_ and FD_below_), and the Community-Weighted Mean (CWM) of each trait.

Species richness was quantified as the total count of recorded species, while total vegetation cover as the cumulative percentage of plant cover within the plot area.

For each trait, the community-weighted mean (CWM) was computed using the “functcomp” function (package FD; Laliberté et al., 2014). To characterize the functional diversity of plant communities, we calculated Rao’s quadratic entropy (RaoQ) as a measure of overall trait dissimilarity. In addition, to capture other key aspects of functional diversity, we also computed functional richness (FRic) and functional evenness (FEve), which represent the extent of trait space occupied by the community and the regularity of species’ trait distribution within that space, respectively (Villéger et al., 2008). Functional diversity indices were measured separately for above and belowground traits. We first generated a multi-trait dissimilarity matrix across all species ("gawdis" function; de Bello et al., 2021) and then used the “dbFD” function (package FD; Laliberté et al., 2014) to quantify above- and belowground functional dissimilarity (FD_above_ and FD_below_), functional richness (FRic_above_ and FRic_below_), and functional evenness (FEve_above_ and FEve_below_). However, to avoid inflating model complexity and potential multicollinearity, we ran separate models for each functional diversity index. As the results obtained with FRic and FEve were qualitatively similar to those based on RaoQ functional dissimilarity (FD), we present only the models including RaoQ in the main text and report the others in the Supporting Information (Appendix S5 and S6).

#### Ecosystem functioning

Ecosystem functions assessed using more than one variable (see Table 1), i.e. water regulation, soil carbon stock, and nutrient cycling were quantified using the averaging approach (Byrnes et al., 2014; Maestre et al., 2012). This method calculates the mean of the Z-score values across the EF variables, providing a straightforward composite measure of an ecosystem function.

### Statistical analysis

In order to reconstruct past ecosystem functions and assess how they changed over time, we used a stepwise approach (Fig. 1). We first assessed BEF relationships in a subset of vegetation plots and estimated the predictive ability of biodiversity for hindcasting ecosystem functioning. Then, we applied the estimated BEF relationships to predict ecosystem functions of a large resurveying vegetation dataset as well as past ecosystem functioning levels based on historical vegetation data. Finally, we applied this method to explore temporal trends in ecosystem functioning under different protection regimes.

#### Predictive models for ecosystem functions

Past erosion control and invasion resistance were quantified directly from historical vegetation records, as these functions are derived from vegetation data. In contrast, past information on biomass production, carbon stock, water regulation, and nutrient cycling was not available for plots outside the BEF subset or for the historical surveys, and therefore required predictions. Thus, using both random forest <u>(randomforest package; Breiman et al.,</u> <u>2018)</u> and linear mixed-effect models (“lmer” function, lme4 package; Bates et al., 2014), we fit current BEF relationships based on the data of the BEF subset. Separate models were fit for each ecosystem function considered as response variable, i.e. biomass production, soil carbon stock, water regulation, and nutrient cycling. In each model, biodiversity metrics and environmental variables were used as predictors. Before conducting the analysis, we assessed multicollinearity among predictors and excluded those with a Variance Inflation Factor (VIF) exceeding 3 (Zuur et al., 2010; see also Appendix S3). The final predictors included SR, total vegetation cover, FD_above_, FD_below_, H_CWM_, SLA_CWM,_ LDMC_CWM_, RDMC_CWM_, RTD_CWM_, sea distance, and MAT. Additionally, in the mixed-effect models, we incorporated the statistical interaction between sea distance and the biodiversity variables to address potential impacts of stress and disturbance on the BEF relationships (La Bella et al., 2024), resulting in the initial estimation of 20 fixed-effects parameters before model selection. For each response variable, we initially fitted mixed-effects models including study sites as a random factor. The significance of random effects was assessed through likelihood ratio tests comparing models with and without the random term. If the random effect was not significant, it was removed, and the simpler linear model was retained to improve parsimony. We applied a model selection and averaging approach using the “dredge” function (MuMIn R package; Barton & Barton, 2023), which ranks all possible models based on their Akaike Information Criterion (AIC), accounting for both model fit (via log-likelihood) and complexity. We then averaged the parameters of all models with ΔAIC < 2 (Burnham & Anderson, 2002; Richards, 2005). Model residuals underwent visual inspection to ensure homoscedasticity and normality (Zuur et al., 2010). Before the analysis, biomass production was log-transformed to improve model fit and reduce skewness in the response variable.

#### Cross-validation

We used leave-one-out cross-validation (LOOCV) to assess the predictive power of the selected models related to supporting ecosystem functions (Yates et al., 2023). It consisted in randomly leaving out from the model training set one plot per run, which was used as testing set to predict the ecosystem function variable starting from the plant community and environmental data. However, since within each site plots where moderately close to each other due to the narrow spatial conformation of Mediterranean coastal dunes, we applied the B-LOOCV proposed by Ploton et al. (2020) to account for spatial autocorrelation. This method leaves out from the training set the testing plot and all its neighbours included within a spatial buffer. We tested several exclusion buffer sizes, with radius ranging between 0 m and 500 m with intervals of 50 m. This spatial extent has been previously advised for Mediterranean costal dunes (Malavasi et al., 2018). The procedure was repeated 1000 times for each ecosystem function model. For each iteration, the model predictive performance was quantified using several goodness-of-prediction statistics: Pearson correlation between observed and predicted values (Pearson *r*); root mean squared error (RMSE); and mean absolute error (MAE). Response variables were scaled and centred (mean = 0 and standard deviation = 1) before the cross validation in order to obtain standardized and comparable measures of the predictive performance (i.e. not influenced by the unit scale). We then calculated the mean and standard deviation of each statistical metric, grouped by buffer size to evaluate if the model predictive accuracy was affected by spatial autocorrelation. The model performance was consistent across sizes of exclusion buffer (Appendix S7), indicating low spatial dependence between training and test sets. Linear models and random forest performance was qualitatively similar (see Table 2 for linear models and Appendix S8 for random forest), thus we kept only mixed-effect models for predictions.

**Table 2.** Predictive models of ecosystem functions. The table shows the goodness-of-prediction of each model evaluated with several statistics, i.e. Pearson *r*, RMSE, and MAE. “Model” column indicates whether the final model after model selection consisted in a linear model (“lm”) or a linear mixed-effect model (“lmer”). Multiple R^2^ and conditional R^2^ are provided for each model. Pearson *r* is the correlation between observed and predicted values. RMSE stand for root-mean-square error while MAE for mean absolute error.

| EFs | Model | Multiple/<br>conditional<br>$R^2$ | Pearson $r$ | RMSE | MAE |
| --- | --- | --- | --- | --- | --- |
| Biomass production | lm | 0.760 | 0.730 | 0.605 | 0.489 |
| Water regulation | lmer | 0.583 | 0.581 | 0.706 | 0.599 |
| Carbon stock | lmer | 0.603 | 0.566 | 0.740 | 0.627 |
| Nutrient cycling | lmer | 0.531 | 0.524 | 0.832 | 0.691 |

We finally selected BEF models that showed a satisfactory predictive ability (e.g. Pearson *r* > 0.5) and used them to predict the ecosystems functions of the whole resurveying dataset in both time periods, i.e. past and present.

#### Temporal changes in ecosystem functioning across different protection regimes

We evaluated differences in temporal changes in ecosystem functioning in communities under different protection regimes using linear mixed-effect models (“lmer” function, lme4 package; Bates et al., 2014). Ecosystem functions were analysed separately and used as separate response variables. Time, protection status, and their interaction were used as fixed effects. The “Time” variable consisted in two levels, i.e. past and present; while protection status had three levels, i.e. national protected areas, Natura 2000 sites, and non-protected areas. Study site and plot identity nested in study site were included as random intercepts to account for the temporal repeated measure in each plot and for the spatial dependence of plots within sites. We performed a Moran’s test to verify for any remaining spatial autocorrelation of the residuals of each model (“moran.test” function, spdep package; Bivand & Wong, 2018), which revealed no spatial autocorrelation (Appendix S9). For each model, residuals were visually inspected to ensure assumptions of homoscedasticity and normality (Zuur et al., 2010). Finally, Tukey’s post hoc tests were used to assess the significance of temporal trends in the three classes of protection (emmeans package; <u>Lenth, 2022)</u>. We consider non-significant changes as an effective conservation outcome of the protection regime. Since species related to invasion resistance or erosion control were not present in all plots, we analysed temporal trends in these two functions only in plots were non-native or dune-building species were recorded in at least one survey period. For invasion resistance and erosion control, which are proportion variables bounded between 0 and 1, we used beta regression models (glmmTMB package; Brooks et al., 2017). Prior to the analysis, we transformed the response variables to avoid extremes of 0 and 1 (response variable value/1× (number of observations − 1) + 0.01)/number of observations; Cribari-Neto & Zeileis, 2010). We then used diagnostic plots of the DHARMa package (Hartig, 2017) and tested the model for uniformity, dispersion, and homoscedasticity. To account for uncertainty in the predicted past ecosystem functions, we propagated uncertainty from fixed effects, random effects, and residual variance. We first obtained 1000 samples of the model’s coefficients using the “sim” function (arm package; <u>Gelman et al., 2015)</u>. For each set of simulated coefficients, we computed the predicted values of the modelled functions and added normally distributed residual noise with standard deviation equal to the model’s residual standard deviation (function “sigma”), to account for unexplained variability. Then, to propagate the uncertainty we fit the above-mentioned mixed-effect models testing temporal change in ecosystem functions across different protection levels for each set of simulated predictions (n = 1000), generating a distribution of results that incorporates the three sources of uncertainty, i.e. from fixed effects, random effects, and residual variance. Finally, we extracted the mean values and assessed the significance of temporal changes for each protection level using confidence intervals (Appendix S10).

To evaluate the robustness of our findings on the effects of protection status on temporal changes in ecosystem functioning, while accounting for external confounding factors (Geldmann et al., 2018; Joppa & Pfaff, 2009), we performed additional analyses having first pre-processed data using propensity score matching (PSM). We performed pairwise comparisons between protection levels. For each comparison, we matched plots based on environmental similarity (MAT, distance from the sea, and latitude) using nearest-neighbour matching with replacement and a 1:1 ratio (function “matchit”, R package MatchIt; Ho et al., 2007). Covariate balance was evaluated using standardised mean differences and Love plots. We then tested for differences in the magnitude of temporal change in ecosystem function (ΔEF = EF_t0_ − EF _t1_) between pairs of protection levels (i.e. "non-protected" vs "Natura 2000", " non-protected " vs "national protected areas", "Natura 2000" vs "national protected areas") using Welch’s t-tests on matched samples, to assess whether the degree of protection influenced the observed changes over time. The results of this analysis confirmed the general pattern observed with the mixed-effect models not accounting for matching (Appendix S11).

## RESULTS

### Predictive models for ecosystem functions

The models of biomass production, water regulation, carbon stock, and nutrient cycling showed a satisfactory predictive ability (Pearson *r* > 0.5; Table 2). Among these, the model for biomass production had the highest predictive ability (Pearson *r* = 0.8).

Across the models, total vegetation cover and H_CWM_ were the most important predictors of ecosystem functioning (Fig. 2a), with a general positive effect across all functions, suggesting that in open, spatially heterogeneous habitats such as coastal dunes, structural dominance and space occupancy may be the most important drivers. Other biodiversity component also contributed to ecosystem functioning, but their effect was less consistent. Communities with low species richness and higher RTD improved biomass production. Communities dominated by species with high SLA and low functional diversity in the aboveground traits supported greater water regulation. Low functional diversity in the aboveground traits promoted soil carbon stock. Finally, we also tested the interaction between biodiversity metrics and sea distance, and we found that some biodiversity effects vary along the sea-inland environmental gradient (Fig. 2b). For example, the effect of aboveground functional diversity on nutrient cycling decreased along the sea-inland gradient.

**Figure 2.**
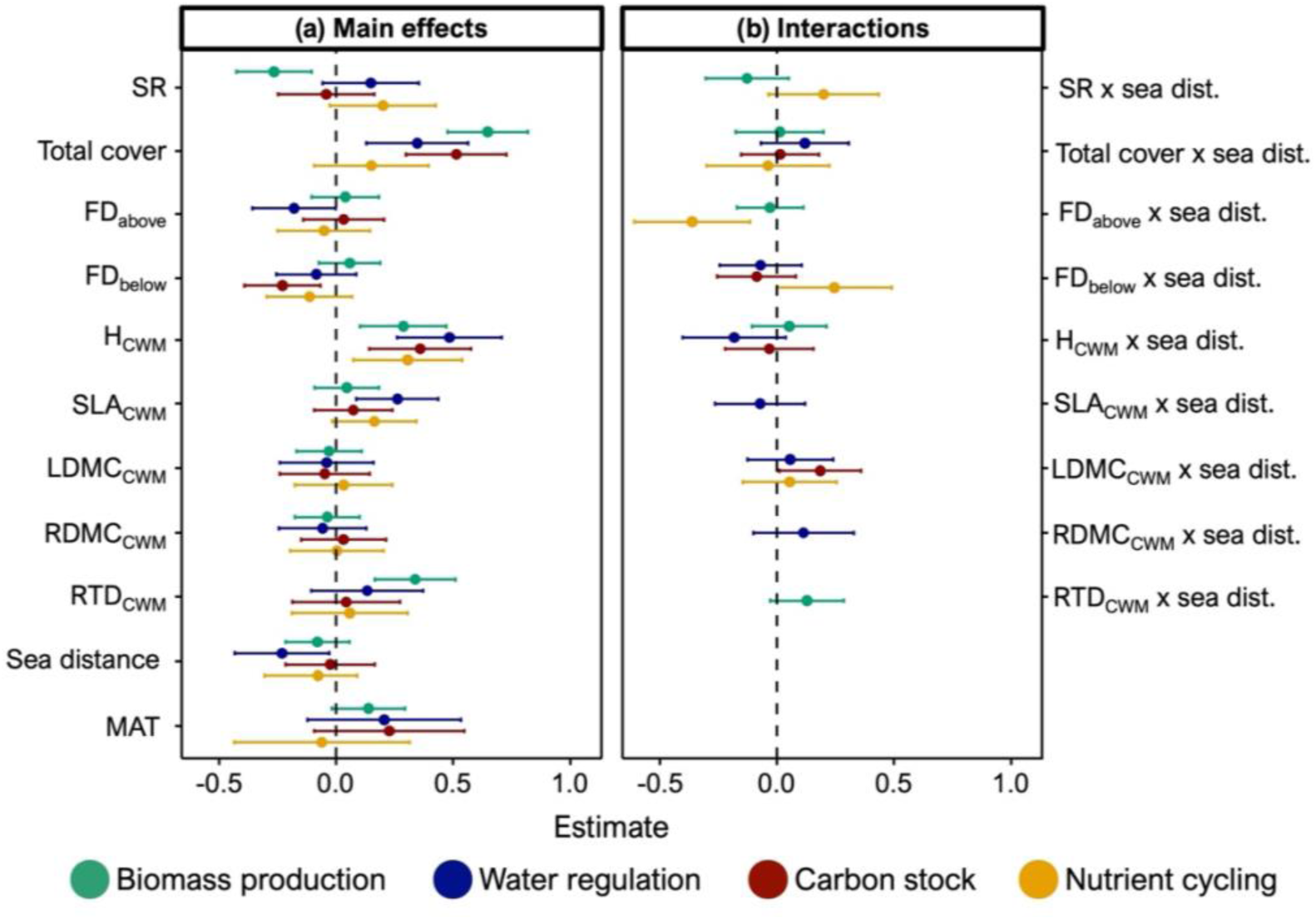
Results of the predictive models. Dots indicate the standardized regression coefficients while error bars the associated 95% confidence intervals. Panel (**a**) shows the direct effects of biodiversity and environmental conditions on ecosystem functioning. Panel (**b**) refers to the interaction coefficients which describe how the effect of the biodiversity metrics on ecosystem functions varies along the sea inland gradient. A negative effect size would indicate e.g. that the effect of SR on EF decreases as sea distance increases. SR, species richness; FD, functional diversity (RaoQ); H, height; SLA, specific leaf area; LDMC, leaf dry matter content; RDMC, root dry matter content; RTD, root tissue density; MAT, mean annual temperature; sea dist., sea distance.

### Temporal changes in ecosystem functioning across different protection regimes

We identified temporal changes in all ecosystem functions. However, these changes were not consistent across different protection regimes. Specifically, in non-protected areas, biomass production, water regulation, and carbon stock increased over time (respectively: estimate = 0.491, *P* = 0.003; estimate = 0.767, *P* = 0.001; estimate = 0.516, *P* < 0.001; Fig. 3a-c) while invasion resistance decreased (estimate = -0.721, *P* < 0.001; Fig. 3f). Inside Natura 2000 sites, water regulation increased (estimate = 0.403, *P* < 0.001; Fig 3b) while erosion control and invasion resistance decreased over time (respectively: estimate = - 0.838, *P* < 0.001; estimate = -0.444, *P* < 0.001; Fig 3e-f). In contrast, within national protected areas most functions remained stable over time and invasion resistance even increased (estimate = 0.722, *P* < 0.001; Fig. 3e).

**Figure 3.**
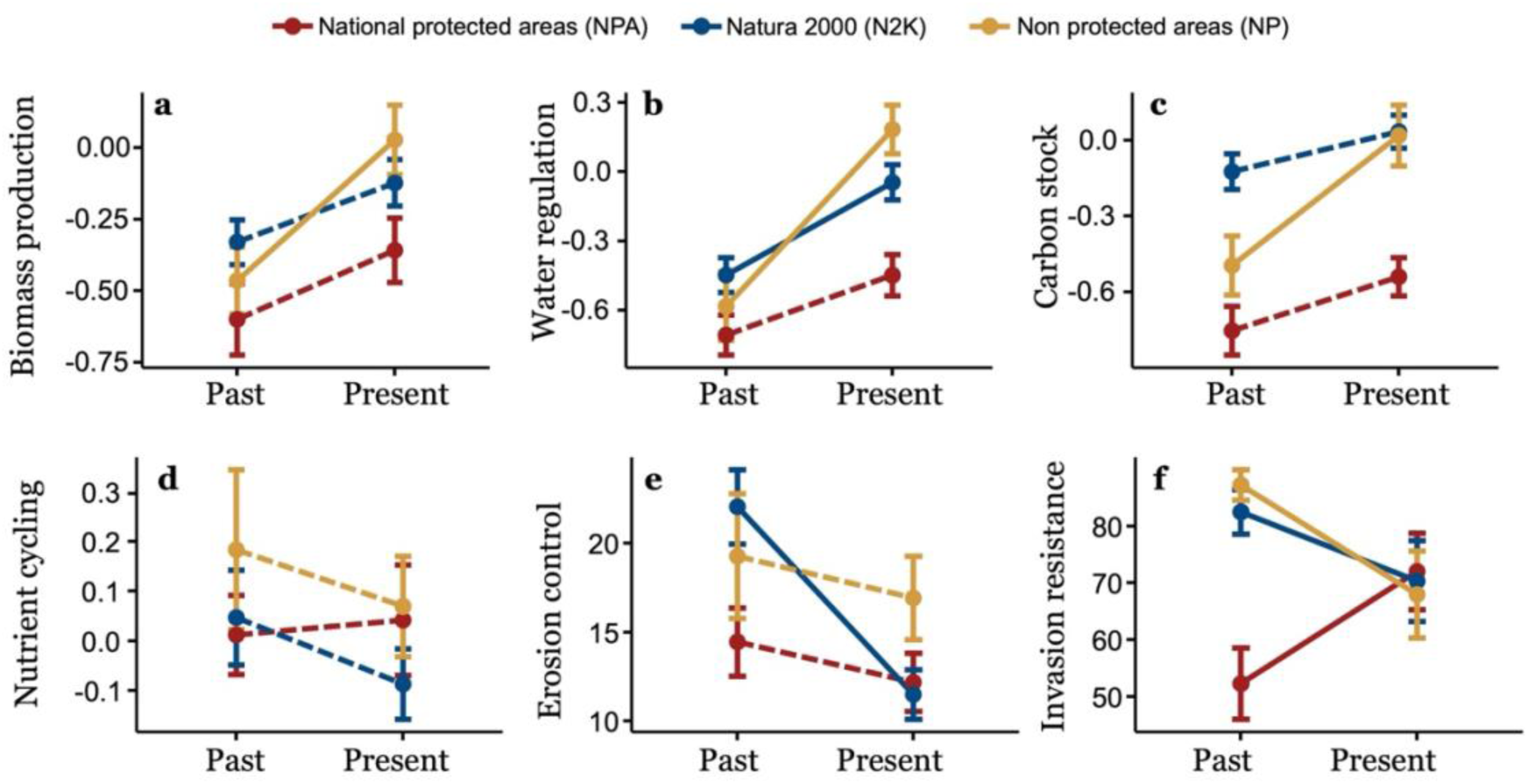
Temporal changes in ecosystem functions. Dots indicate the mean values and lines the associated standard errors. Continuous lines refer to significant changes, while dotted lines to non-significant.

## DISCUSSION

In this study, we proposed a new application of revisitation studies in combination with locally estimated BEF relationships to trace back past ecosystem functions and assess how these have changed over the last 10-15 years under different regimes of protection. Specifically, in this section, we first discuss the ability of biodiversity to predict ecosystem functioning in coastal dunes as well as the effectiveness and limitations of using BEF relationships to hindcast past ecosystem functioning. Then, we highlight the relevance for conservation of our approach by proposing a possible implementation, i.e. monitoring how ecosystem functioning has changed in the context of ongoing global changes as well as the performance of different protection regimes in counteracting these temporal changes in the study area.

### Biodiversity as predictor of past ecosystem functioning

In this study, we tested the ability of plant diversity and community composition to model several ecosystem functions in coastal dunes, such as biomass production, water regulation, carbon stocks, and nutrient cycling. Since BEF relationships are often highly context dependent (Jucker et al., 2016; La Bella et al., 2024; Ratcliffe et al., 2017; Spohn et al., 2023), our models also include environmental conditions and how these interact with the effect of plant diversity and community composition on ecosystem functioning. Including the environmental context is necessary to properly model BEF relationships, and it improves the predictive power of biodiversity (Díaz et al., 2007; La Bella et al., 2023, 2024; Lavorel et al., 2011; Van Der Plas et al., 2020). Here, we found that plant diversity, community composition, and environmental conditions can effectively predict the ecosystem functions analysed. Among all functions, biomass production was the one that was best predicted. This result is in line with the strong plant diversity-biomass relationship found in a wide range of natural ecosystems (Duffy et al., 2017; Garnier et al., 2015; van der Plas, 2019). In contrast, the predictive ability of the models for carbon stock, water regulation, and nutrient cycling was lower compared to biomass production, but still sufficient to make meaningful predictions.

Interestingly, we found that some ecosystem functions tend to decrease along the sea-inland gradient, indicating that close to the sea ecosystem functions are generally higher. This pattern is in line with previous studies showing that in coastal dunes some ecosystem functions, such as pollination, can display strong spatial heterogeneity, with functional peaks often associated with pioneer communities near the seashore (e.g., drift lines and shifting dunes), due to the presence of keystone species (Fantinato et al., 2018).

### Limitations of hindcasting ecosystem functions

While our study highlights the potential of using revisitation data and BEF relationships for hindcasting past ecosystem functions, some limitations should be acknowledged. First, the availability and quality of historical vegetation data pose a constraint, as they can vary in taxonomic resolution, plot re-location accuracy, and sampling methodologies, potentially introducing biases in reconstructing past ecosystem functioning. Second, our hindcasting approach assumes that relationships between biodiversity, environmental conditions, and ecosystem functions remain stable over time. This is a common assumption in predictive ecological models including forecasting (Dietze, 2017), species distribution models (Elith & Leathwick, 2009) and trait-based projections of community or ecosystem change (Laughlin & Messier, 2015). However, we acknowledge that BEF relationships may shift due to changes in environmental drivers, disturbance regimes, or species interactions. While we assessed spatial model accuracy via cross-validation, we could not evaluate temporal predictive performance due to a lack of independent historical data. To overcome these two issues, and to minimise uncertainty and avoid extrapolation errors, we restricted predictions to a subset of the original revisitation dataset, excluding 30% of plots. Specifically, we excluded plots belonging to highly variable or ephemeral habitats (namely the drift lines’ annual vegetation), plots lost due to anthropogenic changes or coastal erosion, and plots with environmental conditions or biodiversity values falling outside the models’ prediction range (see Methods for details). Moreover, our simulation-based approach incorporates parameter and residual uncertainty, improving robustness.

Although our hindcasting framework has inherent limitations, it can provide valuable insights where long-term ecosystem function data are lacking. By reconstructing past ecosystem functions based on historical vegetation records, it offers a novel tool to assess long-term ecological changes and support conservation planning. While further testing across ecosystems is needed, this approach can be particularly relevant in light of the growing availability of resurvey data, such as ReSurveyEurope (Knollová et al., 2024), ReSurveyDunes (Acosta et al., 2025), BioTIME (Dornelas et al., 2025), LOTVS (Sperandii et al., 2022), or VESTA (Świerkosz & Reczyńska, 2022).

### Temporal changes in ecosystem functioning across different protection regimes

One of the main arguments for biodiversity conservation relies on the ability of biodiversity to support ecosystem functioning (Balvanera et al., 2006). However, although many BEF studies have reported positive relationships between biodiversity and ecosystem functioning (Le Bagousse-Pinguet et al., 2019; Maestre et al., 2012), several others have highlighted that these relationships are not universally positive and can vary depending on context (Hagan et al., 2021; van der Plas, 2019). In line with this, our findings show that biodiversity can have positive, negative or no effects on ecosystem functioning. Conserving biodiversity may therefore not necessarily ensure the conservation of ecosystem functioning (O’Connor et al., 2021). To explore this further, we applied our hindcasting approach to compare temporal changes in ecosystem functions across three protection regimes, i.e. national protected areas, Natura 2000 sites, and non-protected areas. Our case study illustrates the utility of our method for conservation and long-term ecological monitoring by revealing how biodiversity changes affect ecosystem functioning.

In the study area, significant vegetation shifts have occurred over the past 10-15 years both inside and outside Natura 2000 sites (Sperandii et al., 2020). As a result, we detected also marked temporal changes in ecosystem functions. Specifically, in non-protected areas and Natura 2000 sites, we observed that most ecosystem functions underwent temporal changes with a general increase of biomass production, carbon stock, and water regulation and a decline in invasion resistance and erosion control. These patterns likely reflect the spread of fast-growing ruderal and non-native species observed in Italian coastal dunes (Del Vecchio et al., 2015, 2016), which often replace native dune-building species responsible of erosion control (Sarmati et al., 2025). Additional trait-based analyses (Appendix S12) supported these trends also in our study area, showing a significant decrease in LDMC_CWM_ in non-protected areas and an increase, although non statistically significant, in SLA_CWM_ in Natura 2000 sites, suggesting a potential shift towards communities with more acquisitive strategies. While ruderal and non-native species may temporarily enhance certain functions, such as biomass production and biogeochemical cycles (Liao et al., 2008; Marzialetti et al., 2025; Rout & Callaway, 2009; Simberloff, 2011; Wilsey et al., 2023), they may, in contrast, undermine long-term ecosystem stability and biodiversity (Del Vecchio et al., 2015; La Bella et al., 2023; Sperandii et al., 2021). It is important to mention that, in coastal dunes, key services include erosion control, invasion resistance, and habitat provisioning for biodiversity (Drius et al., 2019a, 2019b; Liquete et al., 2013), while supporting services such as biomass production, water regulation, and nutrient cycling are relatively low compared to other habitats (Van Der Biest et al., 2017). As such, in these ecosystems higher functioning may not necessarily translate into more valuable services to human societies, and only key functions should be prioritised by conservation actions. Our findings indicate that non-protected coastal dunes, which are not subject to formal management, are shifting toward more productive ecosystems, likely due to the spread of non-native species, with the risk of losing their key characteristics in the near future. On the contrary, Natura 2000 sites are not effectively protecting dune-building species responsible for dune stability, resulting thereby in the loss of erosion control potential, and not contrasting the expansion of non-native species. These findings question the efficacy of the existing set of Natura 2000 sites of our study area to counteract pressures such as tourism, urban expansion, or climate change-driven disturbances, as observed also by other studies (Bricca et al., 2024; Ricci et al., 2023; Tsiafouli et al., 2013) and to prevent the decline of ecosystem functioning or, as showed by Sperandii et al. (2020), of biodiversity.

In contrast, within national protected areas most ecosystem functions remained stable over time and invasion resistance even increased. Although we lack specific data on local management actions, the observed reduction in non-native species suggests the possible influence of control efforts. These results are encouraging, indicating that the national protected areas considered here have been effectively managed to maintain functioning ecosystems beyond biodiversity, even in vulnerable ecosystems like coastal dunes. Our findings support previous research suggesting that stricter protection can enhance ecosystem functioning (Lecina-Diaz et al., 2019), although evidence remains mixed (Castro et al., 2015; Olmo et al., 2025).

The contrasting outcomes between national protected areas and Natura 2000 network likely reflect their distinct objectives and management practices. Natura 2000 network aims to maintain endangered species and habitats at “favourable conservation status” (Commission, 2009), while national protected areas often pursue broader goals, including the conservation of ecosystems and natural processes (CBD, 2022). Still, the most substantial differences lie in management: national protected areas typically benefit from dedicated funding and personnel, whereas Natura 2000 sites often do not (Hochkirch et al., 2013; Kati et al., 2015; Maes et al., 2013). In coastal dunes, effective conservation measures include fencing or wooden walkways to reduce trampling (Acosta et al., 2013; Prisco et al., 2021), invasive species control (Lazzaro et al., 2020), and limitation on mechanical beach cleaning (Battisti et al., 2023; Roig et al., 2009), therefore practices that require considerable financial resources for implementation and long-term maintenance..

Finally, it is important to remember that this work is not intended to assess the overall success of different protection regimes but rather to examine how revisitation data can be used to monitor changes in ecosystem functioning within an existing network of protected areas. Here, we provide insights into the dynamics of ecosystem functioning, highlighting the potential of the hindcasting approach proposed for monitoring and conservation purposes. However, as the number of sites included in this study is relatively limited, the findings from these specific locations should be generalized with caution.

### Final remarks

Although most revisitation studies primarily focus on quantifying biodiversity changes in response to global changes, they also offer a unique opportunity to evaluate temporal trends in the ecosystem functions that biodiversity supports. A major limitation, however, is the lack of direct historical measurements of ecosystem processes. Here, to overcome this data gap, we proposed an innovative practical solution, namely hindcasting past ecosystem functions using present-day BEF relationships. This approach enables to jointly assess temporal changes in biodiversity and ecosystem functioning, providing a more comprehensive understanding of the impacts of global changes on natural ecosystems. In this light, we demonstrated a possible application and the relevance for conservation of this approach, showing how ecosystem functioning has changed over the last 15 years in protected and unprotected coastal dunes.

Many conservation efforts currently focus on preserving individual species or habitats, often neglecting crucial functions and services that ecosystems provide, despite their central role goals and targets of the post-2020 global biodiversity framework. Here, we showed that areas designed to conserve biodiversity can also help maintain ecosystem functions. Still, the mere designation of a protected area does not guarantee conservation success, as evidenced by the functioning decline observed in the Natura 2000 sites of this study. Preserving well-functioning coastal dune ecosystems may instead require rigorous, well-funded conservation measures, like those implemented in national protected areas. These findings highlight the need to monitor within protected areas not only biodiversity but also trends in ecosystems functioning to ensure long-term conservation success.

## ARTICLE IMPACT STATEMENT

Hindcasting ecosystem functions from historical vegetation data enables monitoring protection efficacy in preserving ecosystem functionality

## AUTHOR CONTRIBUTIONS

GLB, ATRA, and MC conceived and designed the study. GLB collected the data, led the statistical analysis, and wrote the first draft of the manuscript. All authors contributed to results interpretation, made significant contributions to the manuscript, and approved it for publication.

## ACKNOWLEDGMENTS

This work was supported by the Grant of Excellence Departments 2018–2022 and the Grant of Excellence Departments 2023–2026, MIUR Italy. GLB, ATRA, MA and MC acknowledge the support of the project “National Biodiversity Future Center – NBFC" funded under the National Recovery and Resilience Plan (NRRP), Mission 4 Component 2 Investment 1.4 - Call for tender No. 3138 of 16 December 2021, funded by the European Union – NextGenerationEU. Project code CN_00000033, Concession Decree No. 1034 of 17 June 2022 adopted by the Italian Ministry of University and Research, CUP F83C22000730006.

