## Supplementary for "Using resurvey data to predict changes in ecosystem functioning across protected and unprotected coastal dunes"

**SUPPORTING INFORMATION**

**Appendix S1** Map showing the study area.

**Appendix S2** Detailed description of study sites and protected areas.

**Appendix S3** Correlation matrix between explanatory variables

**Appendix S4** Ecosystem function variables: sampling and soil analysis.

**Appendix S5** Predictive models with FRic

**Appendix S6** Predictive models with FEve

**Appendix S7** Model performance for linear and random forest models across different sizes of exclusion buffer.

**Appendix S8** Predictive models of ecosystem functions fitted with random forest.

**Appendix S9** Moran’s test results

**Appendix S10** Mean estimate and confidence intervals of the temporal changes in ecosystem functions occurred within each protection level after uncertainty propagation.

**Appendix S11** Temporal changes in ecosystem functioning after propensity score matching (PSM) for protection status.

**Appendix S12** Temporal changes in community weighted mean values.

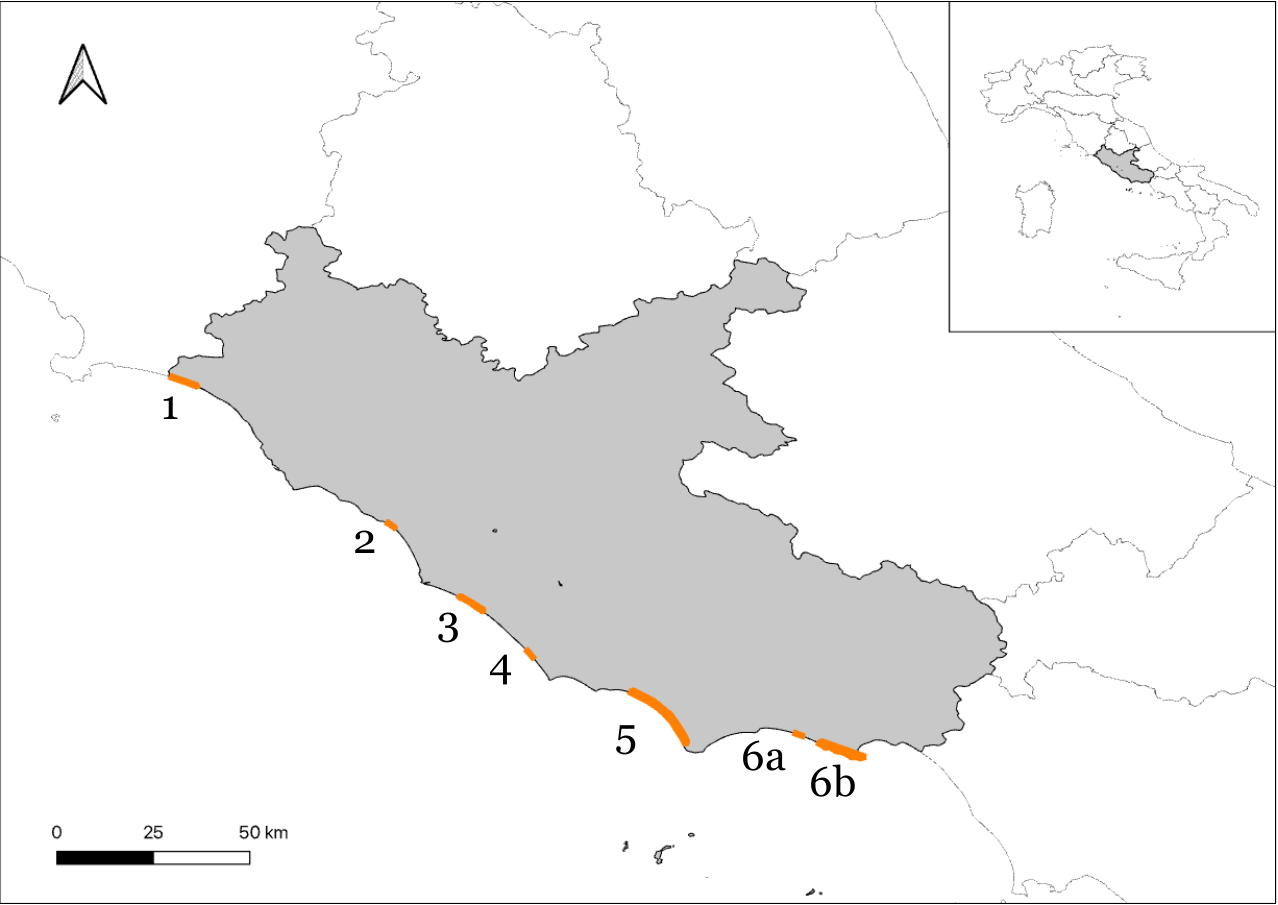

**Appendix S1** Map showing the study area. Numbers refer to study sites (for further details about protected areas see Table S1). 1, Montalto di Castro; 2, Fiumicino; 3, Ostia; 4, Tor San Lorenzo; 5, Sabaudia; 6, Sperlonga.

**Appendix S2** Detailed description of study sites and protected areas. ID numbers refer to Appendix S1. PA, protected area; SAC, Special Area of Conservation; SPA, Special Protection Area.

| **ID** | **Site** | **PA Name** | **Protection type** | **Designation** | **Establishment** |
| --- | --- | --- | --- | --- | --- |
| 1 | Montalto di Castro | IT6010018 - Litorale a NW delle foci del Fiora | Natura 2000 | SAC | 1995 |
| 2 | Fiumicino | Riserva Naturale del Litorale Romano | National PA | State Nature Reserve | 1999 |
| 3 | Ostia | IT6030084 - Castel Porziano | Natura 2000 | SPA, SAC | 1998 |
| 4 | Tor San Lorenzo | IT6030045 - Lido dei Gigli | Natura 2000 | SAC | 1995 |
| 5 | Sabaudia | Parco Nazionale del Circeo | National PA | National Park | 1934 |
| 6a | Sperlonga | IT6040021 - Duna di Capratica | Natura 2000 | SAC | 1995 |
| 6b | Sperlonga | IT6040022 - Costa rocciosa tra Sperlonga e Gaeta | Natura 2000 | SPA | 1995 |

**Appendix S3** Correlation matrix between explanatory variables

|  | **SR** | **Total cover** | **H_CWM_** | **LA_CWM_** | **SLA_CWM_** | **LDMC_CWM_** | **RD_CWM_** | **BODMC_CWM_** | **RTD_CWM_** | **SRL_CWM_** | **FD_above_** | **FD_below_** | **FRic_above_** | **FRic_below_** | **FEve_above_** | **FEve_below_** | **MAP** | **MAT** | **Bio18** | **Bio11** |
| --- | --- | --- | --- | --- | --- | --- | --- | --- | --- | --- | --- | --- | --- | --- | --- | --- | --- | --- | --- | --- |
| **Sea dist** | 0.31 | 0.26 | -0.22 | -0.17 | 0.24 | 0.39 | -0.21 | -0.01 | 0.14 | 0.19 | 0.02 | -0.03 | 0.09 | 0.20 | 0.08 | 0.07 | -0.12 | -0.01 | 0.09 | 0.07 |
| **SR** | 1.00 | **0.76** | -0.07 | 0.10 | 0.26 | 0.22 | -0.16 | -0.14 | 0.32 | -0.01 | 0.37 | 0.29 | 0.37 | 0.47 | 0.17 | 0.10 | -0.25 | -0.23 | -0.28 | -0.16 |
| **Total cover** |  | 1.00 | 0.08 | 0.24 | 0.27 | 0.15 | -0.19 | -0.03 | 0.45 | -0.10 | 0.46 | 0.37 | 0.38 | 0.43 | 0.19 | 0.06 | -0.26 | -0.23 | -0.26 | -0.18 |
| **H_CWM_** |  |  | 1.00 | **0.81** | -0.67 | -0.24 | 0.25 | 0.13 | 0.10 | -0.43 | 0.37 | 0.25 | 0.06 | 0.24 | 0.03 | 0.02 | -0.01 | 0.01 | -0.01 | 0.02 |
| **LA_CWM_** |  |  |  | 1.00 | **-0.71** | -0.42 | 0.45 | -0.17 | 0.22 | -0.58 | 0.39 | 0.44 | 0.20 | 0.31 | 0.07 | -0.16 | 0.03 | 0.06 | 0.03 | 0.09 |
| **SLA_CWM_** |  |  |  |  | 1.00 | 0.22 | -0.28 | -0.07 | 0.12 | 0.33 | -0.13 | -0.03 | 0.18 | -0.03 | -0.02 | -0.03 | -0.12 | -0.17 | -0.16 | -0.18 |
| **LDMC_CWM_** |  |  |  |  |  | 1.00 | -0.34 | 0.16 | -0.25 | 0.63 | -0.24 | -0.22 | 0.04 | -0.06 | 0.11 | 0.19 | -0.12 | -0.07 | -0.09 | -0.05 |
| **RD_CWM_** |  |  |  |  |  |  | 1.00 | -0.21 | -0.15 | **-0.75** | -0.15 | 0.35 | 0.18 | 0.16 | -0.16 | -0.21 | 0.17 | 0.21 | 0.19 | 0.21 |
| **BODMC_CWM_** |  |  |  |  |  |  |  | 1.00 | 0.36 | 0.08 | 0.14 | -0.01 | -0.16 | -0.19 | 0.10 | 0.23 | 0.23 | 0.24 | 0.19 | 0.19 |
| **RTD_CWM_** |  |  |  |  |  |  |  |  | 1.00 | -0.48 | 0.58 | 0.49 | 0.17 | 0.28 | 0.23 | 0.17 | -0.03 | -0.05 | -0.12 | -0.03 |
| **SRL_CWM_** |  |  |  |  |  |  |  |  |  | 1.00 | -0.40 | -0.58 | -0.23 | -0.37 | 0.01 | 0.07 | 0.06 | 0.01 | 0.07 | -0.06 |
| **FD_above_** |  |  |  |  |  |  |  |  |  |  | 1.00 | 0.63 | 0.39 | 0.69 | 0.11 | 0.16 | -0.10 | -0.09 | -0.14 | -0.03 |
| **FD_below_** |  |  |  |  |  |  |  |  |  |  |  | 1.00 | 0.58 | 0.59 | 0.17 | 0.06 | -0.13 | -0.13 | -0.21 | -0.09 |
| **FRic_above_** |  |  |  |  |  |  |  |  |  |  |  |  | 1.00 | 0.65 | 0.04 | -0.12 | -0.23 | -0.21 | -0.23 | -0.17 |
| **FRic_below_** |  |  |  |  |  |  |  |  |  |  |  |  |  | 1.00 | -0.03 | -0.08 | -0.19 | -0.10 | -0.09 | -0.02 |
| **FEve_above_** |  |  |  |  |  |  |  |  |  |  |  |  |  |  | 1.00 | 0.30 | -0.05 | -0.08 | -0.11 | -0.08 |
| **FEve_below_** |  |  |  |  |  |  |  |  |  |  |  |  |  |  |  | 1.00 | -0.12 | -0.11 | -0.16 | -0.07 |
| **MAP** |  |  |  |  |  |  |  |  |  |  |  |  |  |  |  |  | 1.00 | **0.93** | **0.85** | **0.82** |
| **MAT** |  |  |  |  |  |  |  |  |  |  |  |  |  |  |  |  |  | 1.00 | **0.93** | **0.96** |
| **Bio18** |  |  |  |  |  |  |  |  |  |  |  |  |  |  |  |  |  |  | 1.00 | **0.88** |

In red are highlighted the variables with high correlation (*r* < - 0.7 or *r* > 0.7). Sea dist, sea distance; SR, species richness; H, plant height; LA, leaf area; SLA, specific leaf area; LDMC, leaf dry matter content; RD, root diameter; RDMC, root dry matter content; RTD, root tissue density; SRL, specific root length; FD, functional diversity (RaoQ); FRic, functional richness; FEve, functional evenness; MAP, mean annual precipitation; MAT, mean annual temperature; Bio18, precipitation in the warmest quarter; Bio11, mean temperature in the coldest quarter.

**Appendix S4** Ecosystem function variables: sampling and soil analysis.

*Aboveground biomass*

Aboveground living biomass was used as an integrative indicator of plant biomass production potential across plots. Although widely used in biodiversity–ecosystem functioning research (Marquard et al., 2009), plant biomass is not a direct proxy of net primary productivity in communities with perennial species. Nevertheless, it provides a consistent comparative metric of aboveground biomass accumulation (Sala and Austin, 2000). Our biomass measure should therefore be interpreted as a snapshot of potential biomass production rather than a dynamic estimate of net primary productivity. Specifically, in each plot, we clipped all living plant biomass (1 cm aboveground) during the peak of the growing season within a 1x1 m quadrant located in the plot centre. Afterwards, samples were oven dried at 70 °C for 48 h and then weighed.

*Soil sampling and analysis*

We collected 0.5 kg of soil (0-10 cm depth) at five different points around the centre of each plot, then air-dried for 1 week and sieved by a 2 mm mesh. Soil samples were analysed to obtain 16 soil variables related to the soil carbon stock, water regulation, and nutrient cycling (Le Bagousse-Pinguet et al., 2019; Garland et al., 2021). Specifically, we analysed: total organic carbon (TOC, %) and total carbon (TC, %) as a measure of soil carbon stock; electric conductivity (EC) and water holding capacity (WHC, %) as a proxy of water regulation; total nitrogen (TN, %), CN ratio, and total content of P, K, Mg, Mn, Zn, Ni, Ca, Cu, Cr, and Fe for macro- and micronutrient cycling.

TOC, TC, and TN were determined with the Elementar Soli TOC Cube elemental analyser (Zethof et al., 2019). EC was measured using a conductivity metre in a 1:5 (w/v) water and soil mixture, and WHC was measured gravimetrically as the percentage of water retained by the soil (Robertson et al., 1999).

Macro- and micronutrients were determined in 2 g of dry soil using Mehlich III method (Mehlich, 1984) and then after nitric-perchloric acid digestion with simultaneous plasma emission spectrophotometer (ICP–OES).

**Appendix S5** Predictive models with FRic

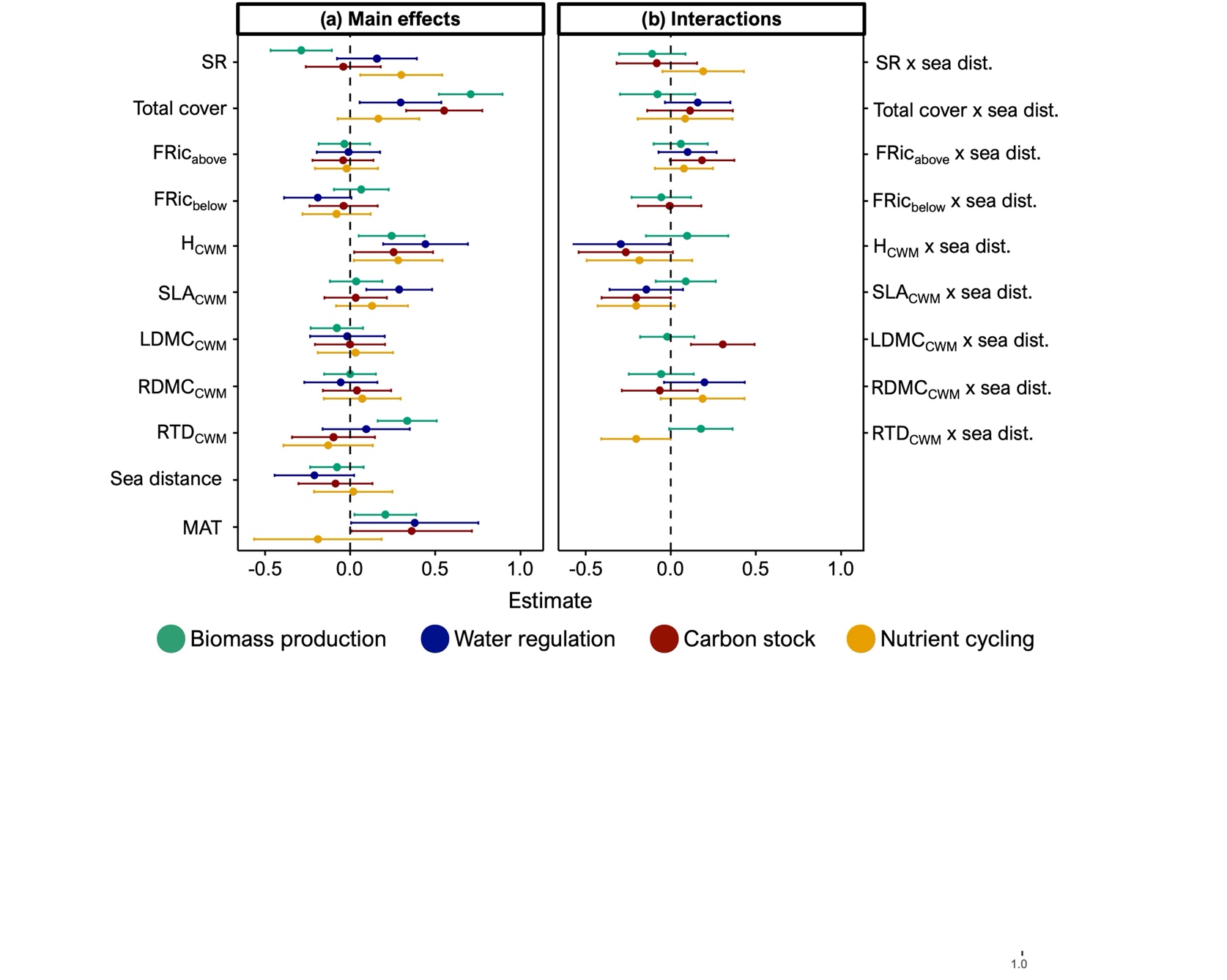

Dots indicate the standardized regression coefficients of each model predictors while lines the associated 95% confidence intervals. Panel (**a**) show the direct effects of biodiversity and environmental conditions on ecosystem functioning. Panel (**b**) refers to the interaction coefficients which describe how the effect of the biodiversity metrics on ecosystem functions varies along the sea inland gradient. A negative effect size would indicate e.g. that the effect of SR on EF decreases as sea distance increases. SR, species richness; FRic, functional richness; H, height; SLA, specific leaf area; LDMC, leaf dry matter content; RDMC, root dry matter content; RTD, root tissue density; MAT, mean annual temperature; sea dist., sea distance.

**Appendix S6** Predictive models with FEve

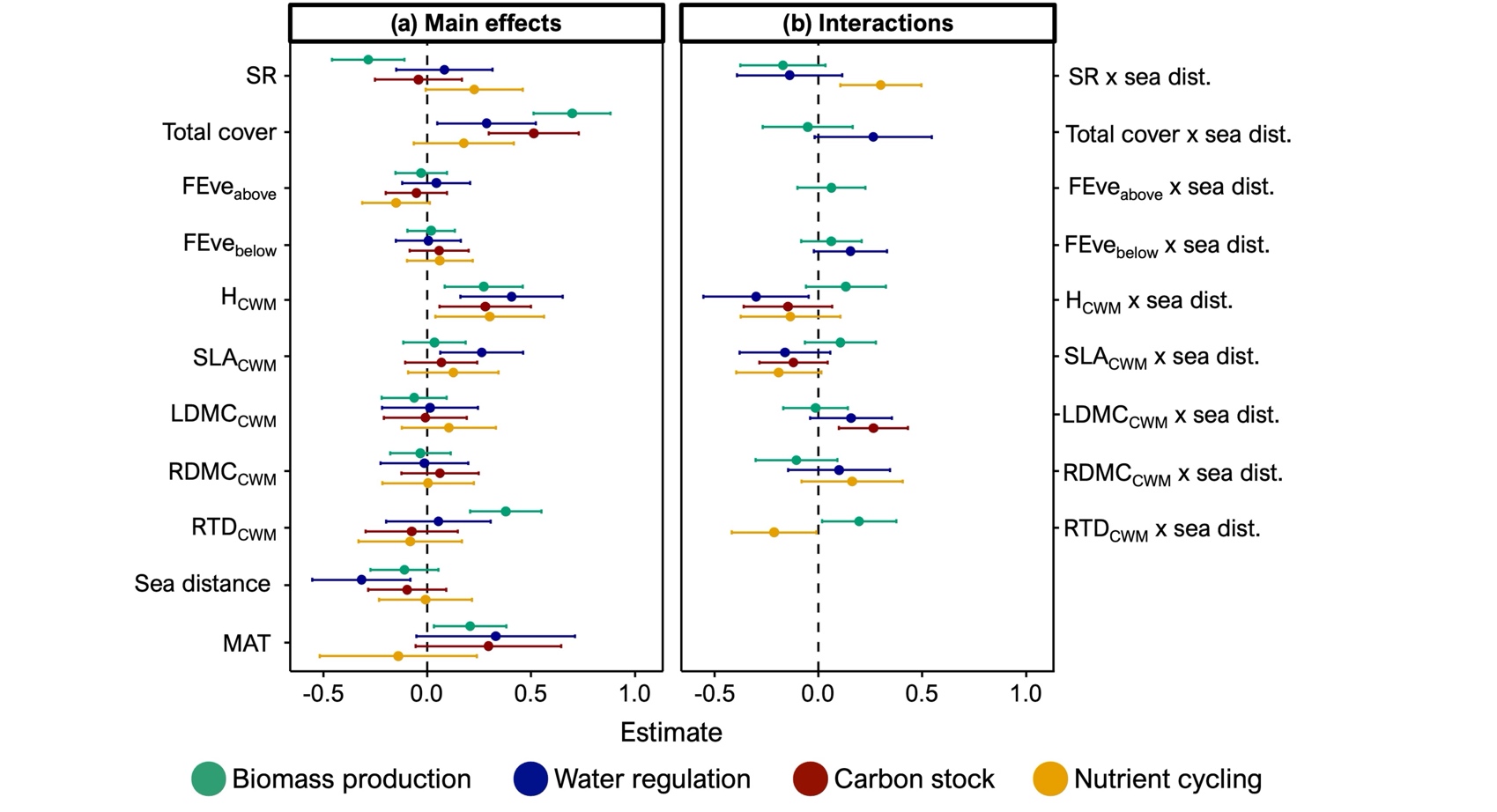

Dots indicate the standardized regression coefficients of each model predictors while lines the associated 95% confidence intervals. Panel (**a**) show the direct effects of biodiversity and environmental conditions on ecosystem functioning. Panel (**b**) refers to the interaction coefficients which describe how the effect of the biodiversity metrics on ecosystem functions varies along the sea inland gradient. A negative effect size would indicate e.g. that the effect of SR on EF decreases as sea distance increases. SR, species richness; FEve, functional evenness; H, height; SLA, specific leaf area; LDMC, leaf dry matter content; RDMC, root dry matter content; RTD, root tissue density; MAT, mean annual temperature; sea dist., sea distance.

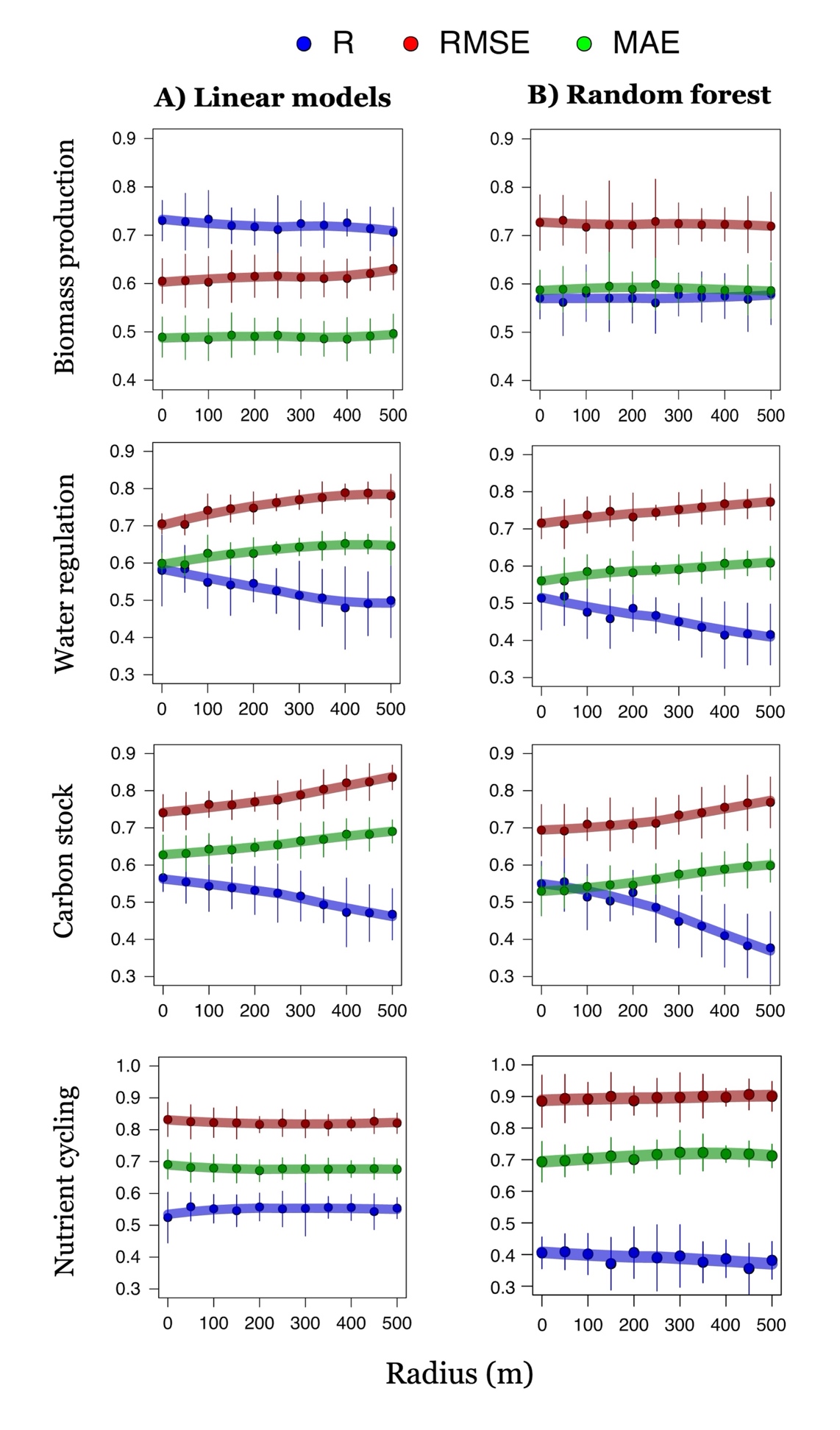

**Appendix S7** Model performance for linear and random forest models across different sizes of exclusion buffer. R indicates the Pearson correlation. RMSE stand for root-mean-square error while MAE for mean absolute error.

**Appendix S8** Predictive models of ecosystem functions fitted with random forest. The table show the goodness-of-prediction of each model evaluated with several statistic: Pearson *r* is the correlation between observed and predicted values; RMSE stand for root-mean-square error; MAE for mean absolute error.

| **EFs** | **Pearson *r*** | **RMSE** | **MAE** |
| --- | --- | --- | --- |
| Biomass production | 0.576 | 0.723 | 0.586 |
| Water regulation | 0.509 | 0.719 | 0.564 |
| Carbon stock | 0.557 | 0.690 | 0.525 |
| Nutrient cycling | 0.407 | 0.883 | 0.693 |

**Appendix S9** Moran’s test results

| **Model** | **Moran I** | ***P*** |
| --- | --- | --- |
| Biomass production | -0.032 | 0.908 |
| Water regulation | -0.032 | 0.912 |
| Carbon stock | -0.031 | 0.907 |
| Nutrient cycling | 0.026 | 0.093 |
| Erosion control | -0.020 | 0.785 |
| Invasion resistance | -0.030 | 0.701 |

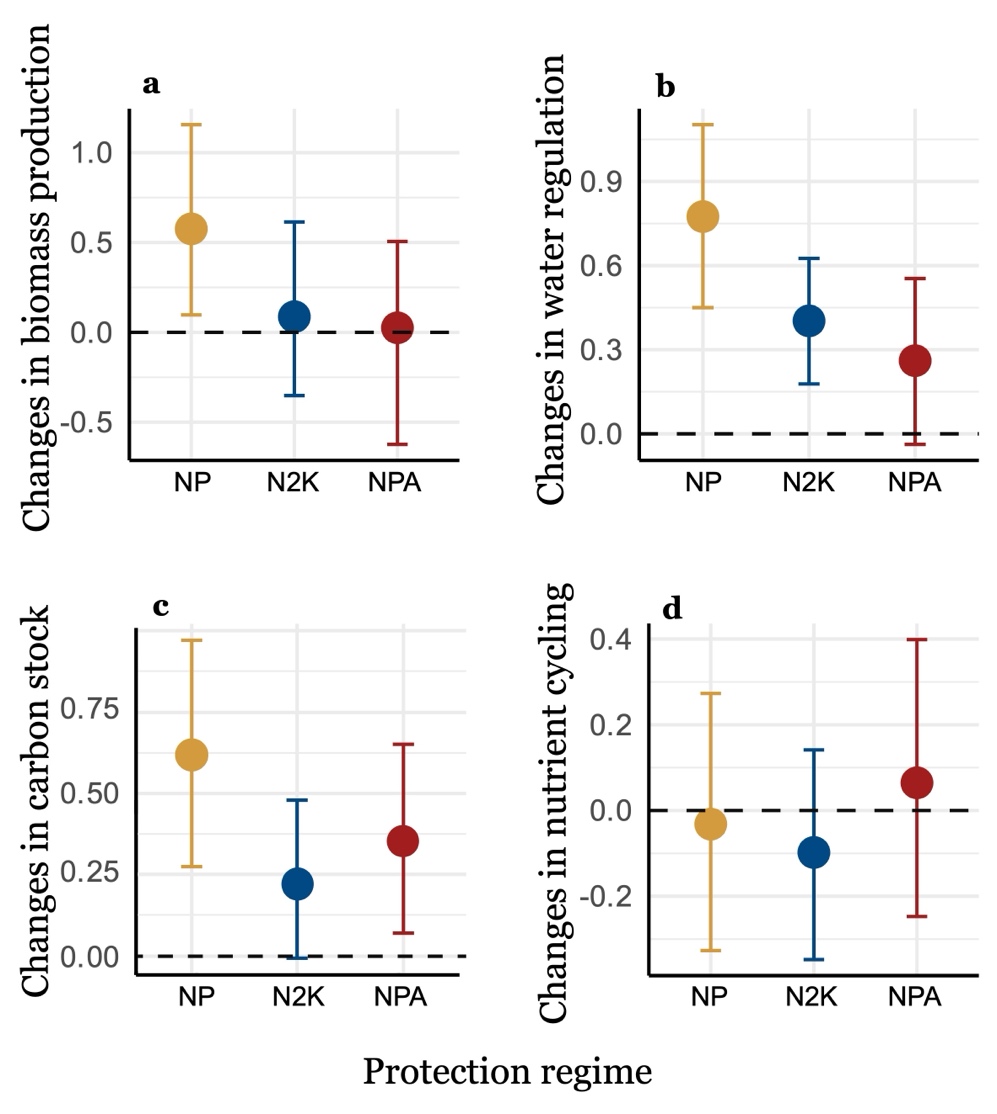

**Appendix S10** Mean estimate and confidence intervals of the temporal changes in ecosystem functions occurred within each protection level after uncertainty propagation.

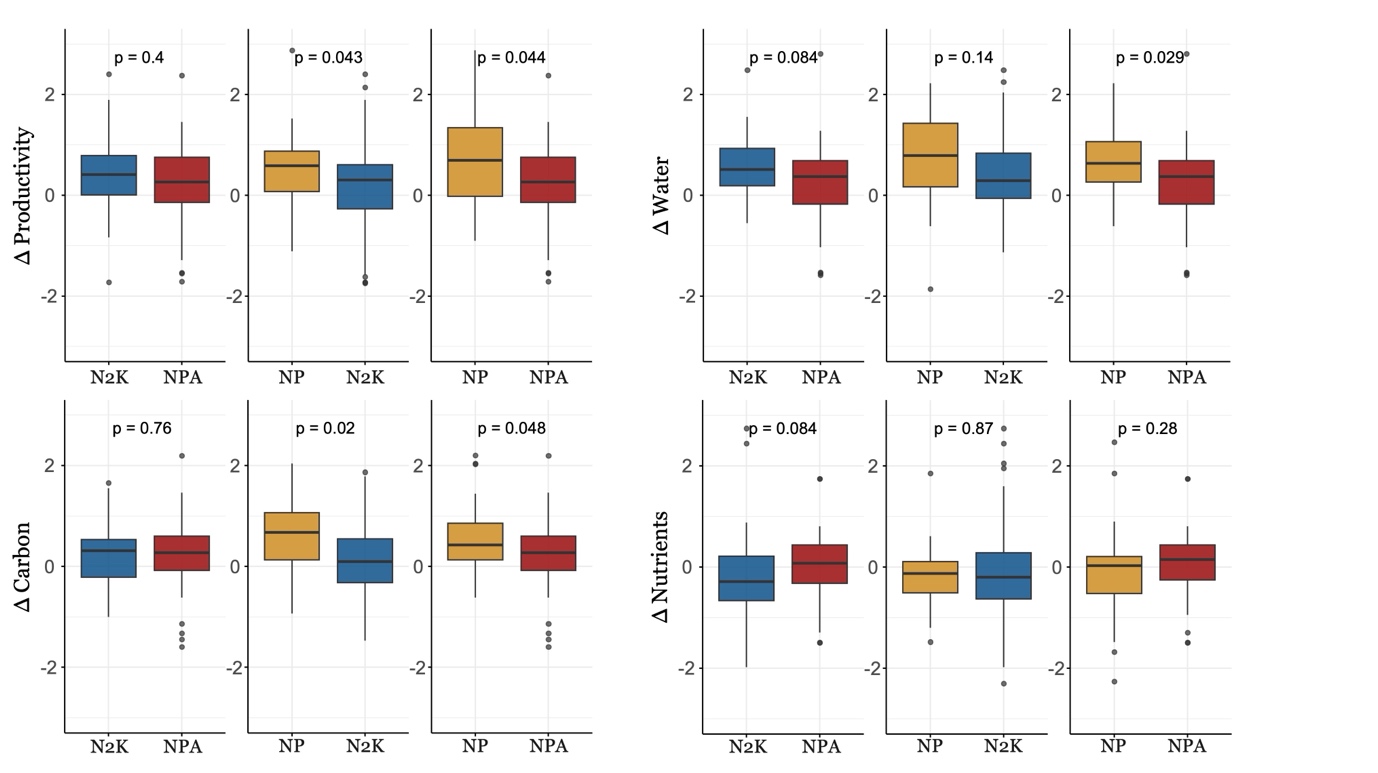

**Appendix S11** Temporal changes in ecosystem functioning after propensity score matching (PSM) for protection status. N2K, Natura 2000; NPA, national protected areas; NP, non-protected areas

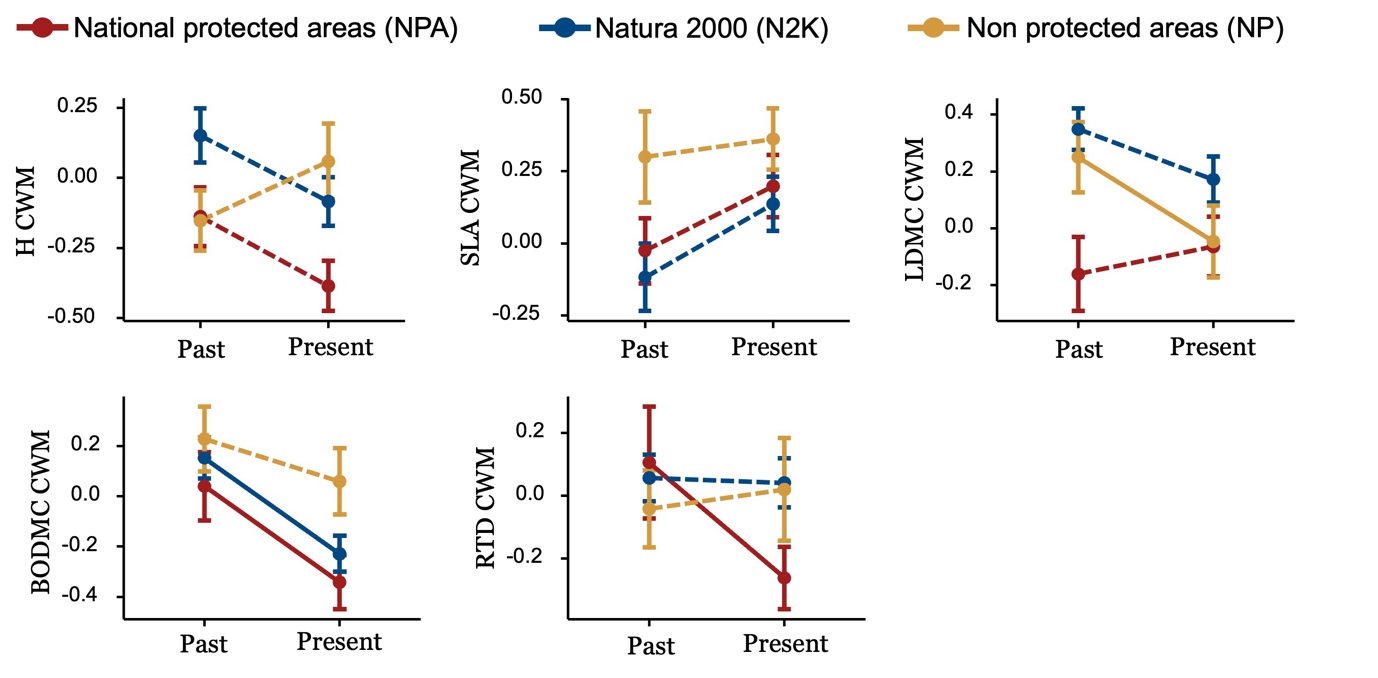

**Appendix S12** Temporal changes in community weighted mean values. Dots indicate the mean values and lines the associated standard errors. Continuous lines refer to significant changes, while dotted lines to non-significant. H, height; SLA, specific leaf area; LDMC, leaf dry matter content; RDMC, root dry matter content; RTD, root tissue density
